# Cornering an Ever-Evolving Coronavirus: TATX-03, a fully human synergistic multi-antibody cocktail targeting the SARS-CoV-2 Spike Protein with *in vivo* efficacy

**DOI:** 10.1101/2021.07.20.452858

**Authors:** Ilse Roodink, Maartje van Erp, Andra Li, Sheila Potter, Sander M J van Duijnhoven, Arthur J Kuipers, Bert Kazemier, Ellen van Geffen, Wieger Hemrika, Bob Berkeveld, Glenn Sonnemans, Britte S de Vries, Bianca Boers, Milou Smits, Sanne Meurs, Maaike de Pooter, Alexandra Thom, Barry N Duplantis, Roland A Romijn, Jeremy Houser, Jennifer Bath, Yasmina N Abdiche

**Affiliations:** ImmunoPrecise Antibodies Ltd

## Abstract

The emergence of severe acute respiratory syndrome coronavirus 2 (SARS-CoV-2) has created an ongoing global human health crisis and will likely become endemic, requiring novel sustainable therapeutic strategies. We report on the discovery of a fully human multi-antibody cocktail (TATX-03) targeting diversified non-overlapping epitopes on the SARS-CoV-2 spike protein that suppressed replication-competent viral titers to undetectable levels in the lungs of SARS-CoV-2 challenged hamsters upon both prophylactic and therapeutic administration. While monotherapy with two of the individual cocktail components also showed clear *in vivo* protection, neither recapitulated the efficacy of TATX-03. This synergistic effect was further supported by examining *in vivo* efficacy of these individual antibodies and corresponding combination therapy at a lower dose. Furthermore, *in vitro* screenings using VSV-particles pseudo-typed with spike proteins representing the SARS-CoV-2 variants of concern Alpha, Beta, and Delta showed that TATX-03 maintained its neutralization potency. These results merit further development of TATX-03 as a potential therapy for SARS-CoV-2 infection with resistance to mutagenic escape.

## INTRODUCTION

The surface glycoprotein (spike protein or S protein) of SARS-CoV-2 is the mediator of host cell attachment and entry and is indispensable for infection^1^. For this reason, most therapeutic monoclonal antibody (Ab) candidates have been directed to the receptor binding domain (RBD) of the S protein to disrupt its interaction with host angiotensin converting enzyme 2 (ACE2) and neutralize host cell entry^2–6^. However, using *in vitro* neutralization as the main predictor for *in vivo* protection ignores other potential mechanisms of action that may occur *in vivo*, including the known contribution of Fc-mediated viral clearance^7^. Additionally, the epitopes of most neutralizing Abs reported in the literature are clustered at or near the ACE2-binding interface of the RBD, leaving many of them similarly vulnerable to escape by evolving viral mutations within the RBD^8^. Use of any therapeutic Ab as a monotherapy significantly increases this risk, necessitating the use of combination therapy approaches to achieve durable protection. However, lack of epitope diversity in the dual-Ab cocktails from Regeneron (casirivimab and imdevimab, targeting adjacent, non-overlapping epitopes) and Eli Lilly (bamlanivimab and etesevimab, targeting overlapping epitopes) leaves both Abs in each cocktail potentially susceptible to evasion by single point mutations^9,10^. Strategies to minimize the risk of viral escape are therefore critical to the effective implementation of Ab-based approaches for SARS-CoV-2 infection prevention and treatment.

Combining three or more monoclonal Abs targeting diversified non-overlapping epitopes on the target has provided transformative prophylactic and therapeutic options for numerous infectious disease targets, including Ebola^11^, respiratory syncytial virus (RSV)^12^, rabies^13^, and botulinum^14^. In addition to minimizing their risk of mutagenic escape, multi-Ab cocktails can stimulate complementary mechanisms of action in concert, unlocking synergistic effects that potentially allow for a lower effective clinical dose than in a monotherapy approach.

Here we describe the discovery of anti-SARS-CoV-2 spike Abs, sourced from phage display panning of our pre-existing human scFv repertoires, reactive to a broad range of epitopes to provide versatile options for multi-Ab combination therapy. By merging neutralization data with epitope binning data, we prioritized TATX-03, a blend of antibodies targeting distinct, non-overlapping epitopes across the spike protein, formulated as two alternate 4-Ab cocktails (TATX-03a and TATX-03b) for *in vitro* and *in vivo* efficacy evaluation. Authentic virus-based *in vitro* neutralization screening revealed that both 4-Ab formulations were able to synergistically neutralize SARS-CoV-2 as only one individual antibody, a component of the TATX-03b cocktail, showed neutralizing effects on its own, but with a clearly higher IC50 compared to the cocktails containing this antibody. Moreover, both formulations provided protection to SARS-CoV-2 infection *in vivo*. We further show that TATX-03 is broadly resistant to emerging variants of concern *in vitro* and we anticipate that it retains *in vivo* efficacy towards these mutants, thereby meriting its further development as a potential therapy for the prevention or treatment of SARS-CoV-2 infection.

## RESULTS

### Phage display library panning enriched for a panel of anti-spike Abs with diverse binding profiles

To enrich for fully human Abs specific for the spike protein, pre-existing human scFv repertoires derived from healthy donors and auto-immune diseased individuals were subjected to four rounds of phage panning against a panel of purified recombinant protein targets using eleven unique panning strategies in parallel **(Fig 1a)**. As determined by ELISA, except for one panning strategy, polyclonal phage outputs showed target-specific binding, with diverse reactivity profiles between the phage outputs from the different panning approaches, suggesting enrichment of phages displaying epitope-diverse Ab fragments. Approximately 700 clones with diverse binding profiles were selected for monoclonal scFv expression and subsequent isolation of periplasmic fractions. Upon confirming their target-specificity by ELISA, the top 279 periplasmic fractions were selected for initial *in vitro* pseudovirus-based neutralization assays. Sixty sequence-unique scFv clones showing diverse reactivity profiles and distinct neutralization capacity were selected for Fv-model based *in silico* developability profiling using BioLuminate (Schrödinger) and subsequent recombinant eukaryotic expression as full-length human IgG1 antibodies. Following Protein A purification and purity/integrity analysis by SDS-PAGE, antibodies were subjected to more in-depth characterization.

**Fig 1.**
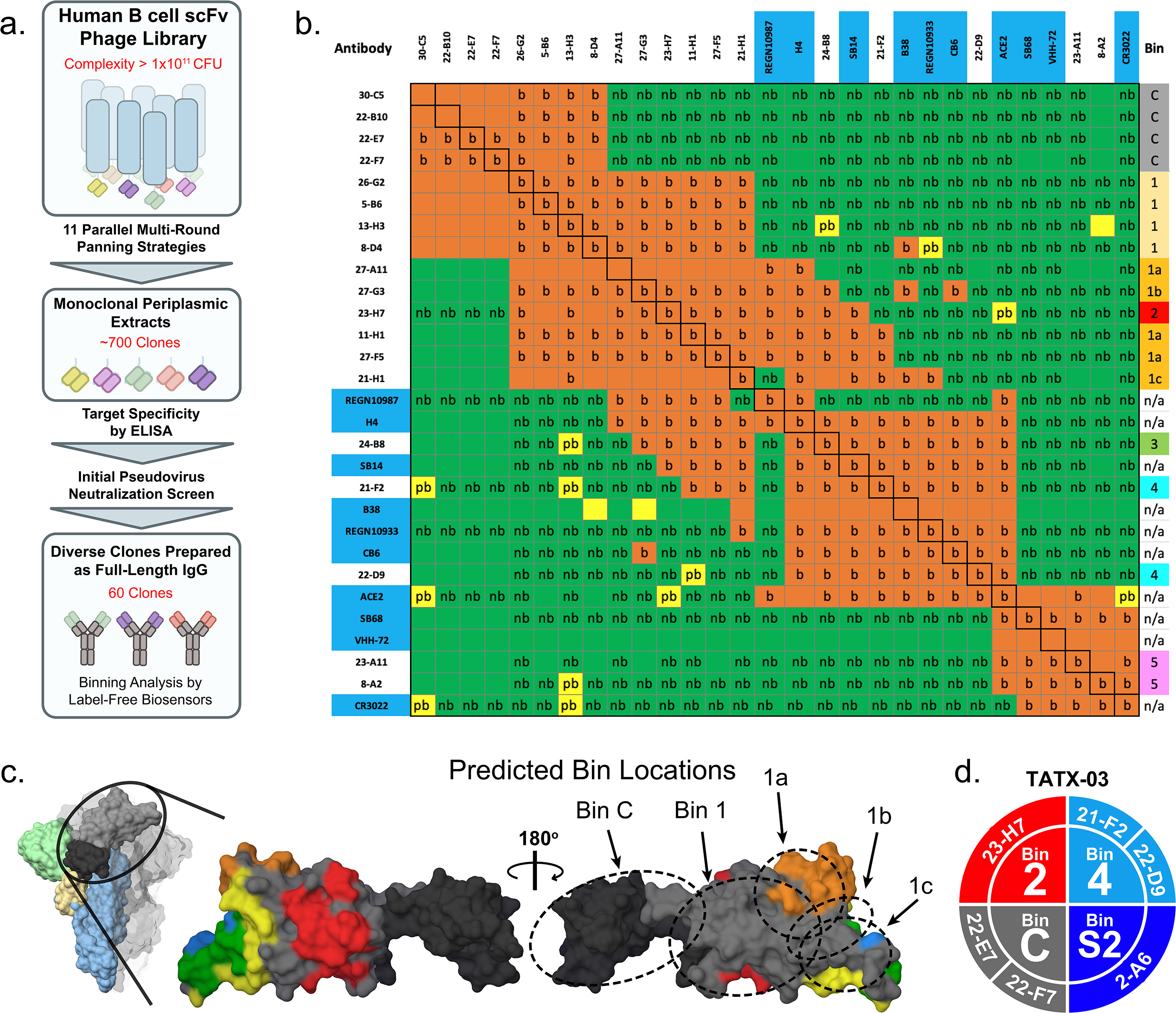
IPA’s library-to-leads triage process. **(a)** High-level schematic of the workflow. **(b)** Heat map for a pairwise analysis of 19 library-derived anti-SARS-CoV-2 S1-specific Abs merged with a panel of ten structural benchmarks (9 literature Abs and ACE2). The red, yellow, and green colored cells indicate Ab pairs that are blocked (b), partially blocked (pb), or not blocked (nb), respectively. Colored cells with a designation of “b, pb, or nb” were measured empirically, whereas those without a designation are “inferred”. The black boxed cells along the diagonal indicate the “self-blocked” pairs. In our bin-definition, bin-members block one another and show similar blocking behaviors when tested against other Abs in the panel. RBD-specific clones were assigned to five bins (1-5). RBD binders that did not block ACE2 were assigned bin “1”, which was split into sub-bins (bin 1a, 1b, and 1c) based on their nuanced blockade towards the structural benchmarks **(Supplemental Fig S1).** A cluster of S1 non-RBD binders blocked bin 1 (but not the sub-bins) and did not block any of the literature Abs, so were assigned to bin “C”, representing the C-terminal nub of the S1 fragment, between the RBD and the furin cleavage site, distinct from the N-terminal domain (NTD); hence, bin C’s specificity is assigned as S1-nonRBD-nonNTD. One RBD-binder Ab (23-H7) blocked bin 1, REGN10987 and uniquely perturbed/partially blocked ACE2 **(Supplemental Fig S2)**, so was assigned to bin 2. Within the ACE2 fully blocking clones, we identified two discrete sets of Abs (bin 4 and bin 5); bin 4 co-located with REGN10933 and CB6, while bin 5 co-located with the “cryptic” epitope of CR3022, VHH-72 and SB68. Bin 3 blocked both bin 2 (23-H7) and bin 4. **(c)** Image of the spike trimer and zoomed in view of the RBD in grey (PDB ID: 7BNM residues 330-520) showing the epitope contacts for benchmark Abs CR3022 (red, PDB ID: 6YMO) REGN10987 (orange, PDB ID: 6XDG), REGN10933 (blue, PDB ID: 6XDG), CB6 (yellow, PDB ID: 7C01), and REGN10933/CB6 shared residues in green. Depiction of the benchmark “bald spot” present on the RBD, an area of the Spike where none of the available literature controls bound. The C terminus of S1-nonRBD is shaded darker grey (residues 320-329; 521-593). Predicted epitope regions for Abs assigned to bin C, bin 1 and sub-bins 1a, 1b, and 1c (dotted ovals) are also indicated. RBD and benchmark antibody structures were imported from PDB to Maestro. Proteins were aligned via Protein Structure Alignment. **(d)** Color wheel showing the composition of TATX-03 blends comprised from six lead Abs distributed across four distinct bins.

### High-throughput epitope binning assays facilitated the identification of multiple distinct epitope bins

An integral part of the triage workflow involved the early implementation of label-free biosensor screenings of the down-selected Abs to assess their pairwise and combinatorial blockade of S-protein by one another, ACE2, and a panel of nine RBD-specific Abs from the literature with known epitopes (REGN10987/imdevimab, REGN10933/casirivimab, CB6/etesevimab, B38, H4, SB14, SB68, VHH-72 and CR3022), as sequences became publicly available. An example heat map resulting from a merged high-throughput binning analysis of our human library-derived clones combined with those from the literature using S1-His(D614G) as target, is shown in **Fig 1b** and highlights the identification of several epitope clusters or “bins” of Abs sharing similar blocking profiles. The inter-bin blocking relationships revealed a series of both overlapping and non-overlapping bins, as shown in the simplified Venn Diagram in **Supplemental Fig S1a**. Literature clones (CR3022, REGN10987, REGN10933, CB6) served as “structural benchmarks” to infer the approximate locations of our deduced bins onto the spike protein, as shown for bin C, bin 1, and its sub-bins a-c **(Fig 1c)**. All S1-non-RBD binders fell into bin C, while the RBD binders were distributed across five bins (1-5). Neither bin C nor bin 1 blocked ACE2 or any of the structural benchmarks. Some bin-1-like clones did not block bin C but showed nuanced interference with binding of some of the benchmarks, so were assigned to sub-bins 1a, 1b, and 1c (**Supplemental Fig S1b-d).**

Bin 2 co-located with REGN10987 and uniquely kinetically perturbed/partially blocked ACE2 **(Supplemental Fig S2a)**. Bin 4 **(Supplemental Fig S2b)** and bin 5 both blocked ACE2 but not one another; bin 4 co-located with REGN10933 and CB6, while bin 5 co-located with the “cryptic” epitope of CR3022, VHH-72 and SB68. Bin 3, like bin 4, also interfered with ACE2 binding and co-located with REGN10933 and CB6, but additionally blocked bin 2, appearing to be a “broader” blocker than bin 4.

While our bin-definition was an over-simplification of a much more nuanced epitope landscape with crosstalk between otherwise discrete bins, it guided our identification of clones from distinct non-overlapping bins that could be curated into cocktails, such as the four-bin combination that formed the basis of our TATX-03 cocktail **(Fig 1d)**.

### Multi-Ab epitope binning experiments confirmed that up to four Abs can co-exist on the spike trimer

Having identified antibody pairs that could co-exist on recombinant monomeric S1-subunit as judged by our pairwise binning matrix, we extended the analysis to higher-order binning experiments using a fully assembled recombinant spike trimer to test whether it could physically accommodate Abs from up to four distinct non-overlapping bins as present in TATX-03 (2, 4, C, and S2). The results from assays performed in complementary formats, a tandem cocktail assay **(Fig 2a and Supplemental Fig S3a,b)** and a premix assay **(Fig 2b and Supplemental Fig S3c-e),** confirmed that Abs from these four bins could access their epitopes without interfering with one another’s binding, thereby validating this bin combination for use in functional studies.

**Fig 2.**
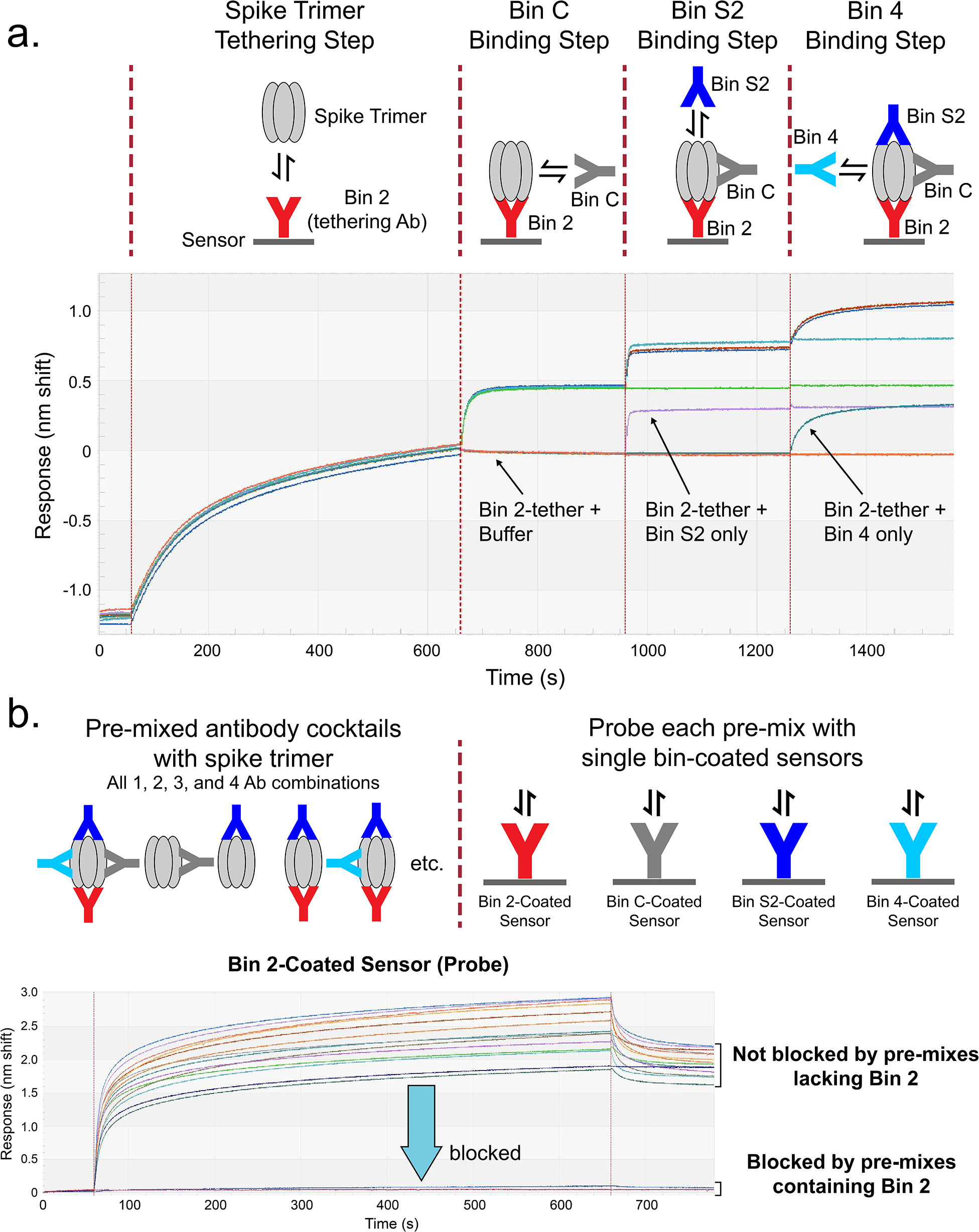
Multi-Ab epitope binning results using complementary assay formats. Binning results verifying the simultaneous saturation of spike trimer with up to four Abs targeting distinct non-overlapping epitopes. **(a)** Tandem cocktail experimental scheme and sensorgrams showing that tethering spike via sensors coated with 23-H7 (bin 2) allows for the stepwise association of Abs from three other non-overlapping bins represented by 22-F7 (bin C), 21-F2 (bin 4), and 2-A6 (bin S2). **(b)** Premix experimental scheme and example sensorgrams. Spike was premixed in solution phase with saturating concentrations of up to four Abs from non-overlapping epitope bins, used individually or as 2, 3-, or 4-Ab cocktails and these premixed samples were presented to Ab-coated sensors (probes), which were blocked in a bin-specific manner. The Abs used for the premix assays were 23-H7 (bin 2), 21-F2 or 22-D9 (bin 4), 22-F7 or 22-E7 (bin C) and 2-A6 (bin S2). See **Supplemental Fig S3** for additional graphs.

While the tandem binning assay had relied upon avid interactions between bivalent full length IgGs and trimeric spike, we investigated the binding affinities of our Abs to recombinant targets under monovalent conditions using complementary assay orientations on the Octet. Most of our lead Abs showed weak affinities with apparent K_D_ values ranging from 0.1 – 1 µM as characterized by square-shaped sensorgrams that were adequately described by an equilibrium analysis. However, clone 23-H7 (bin 2) uniquely bound RBD (or S1) with a high affinity, giving an apparent K_D_ value of approximately 5 nM, regardless of the assay orientation used (**Supplemental Fig S4 and Supplemental Table S1).**

### Some Ab combinations show synergistic neutralization in vitro

Twenty candidate Abs which had been assigned to epitope bins were subsequently tested individually and as 2-, 3-, 4-, and 5-Ab cocktails in a cell-based pseudovirus neutralization assay using a mini-checkerboard format. We reduced the number of combinations screened by first pairing Abs across bins, identifying synergistic pairs and using those to anchor higher-order cocktails. Our TATX-03 four-bin combination, represented by six leads (23-H7 from bin 2, 22-D9 or 21-F2 from bin 4, 22-E7 or 22-F7 from bin C, and 2-A6 from bin S2) which showed varying neutralization capacity individually in the pseudovirus assay **(Fig 3a),** demonstrated synergistic activity in various multi-Ab combinations (**Fig 3b**). In authentic virus-based neutralization assays, all individual Abs, except 21-F2, showed a lack of neutralization within the assay window when tested at a top concentration of 100 µg/ml (**Fig 3c**). However, multiple 4- and 5-Ab combinations showed potent neutralization at the same total antibody concentration, indicating that synergistic effects improved neutralization of authentic virus (**Fig 3c)**. These *in vitro* data resulted in the prioritization of two 4 Ab-cocktails, number 1 (TATX-03a) and 2 (TATX-03b), for *in vivo* efficacy evaluation.

**Fig 3:**
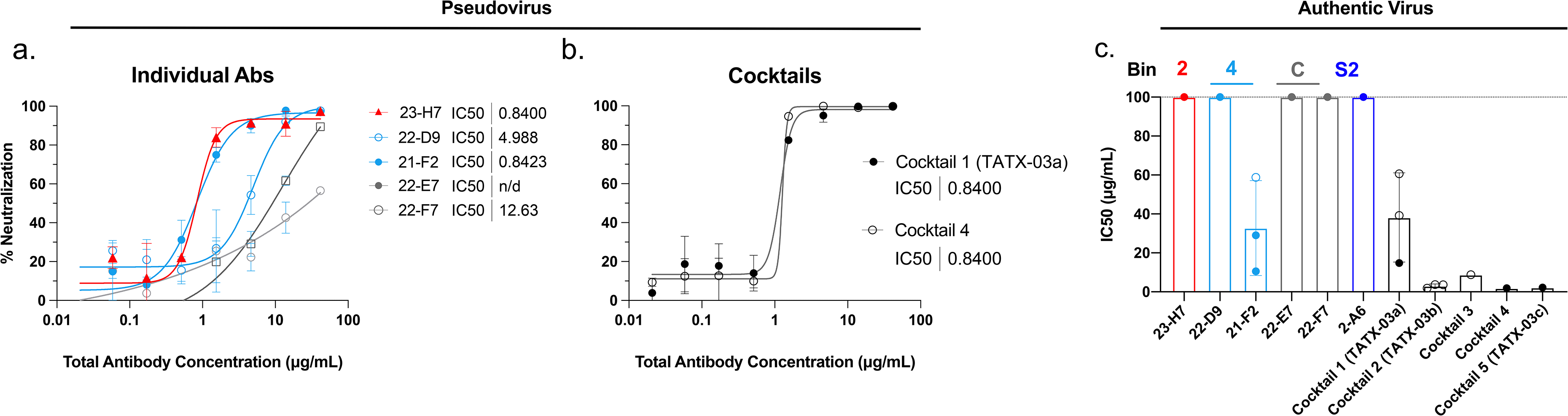
In *vitro* cell-based neutralization data for representative multi-Ab cocktails comprising Abs from bins 2, 4, C, S2. **(a)** Pseudovirus (Wuhan-1 isolate spike sequence) neutralization by individual clones in bins 2, 4 and C respectively, with IC50 values reported. 2-A6 (bin S2) was non-neutralizing in this assay. (**b)** Pseudovirus neutralization of two 4-Ab cocktails with IC50 values reported. **(c)** IC50 values of single Abs and cocktail combinations in authentic virus (D614G strain) neutralization assays. Dotted line represents non-neutralizing limit of detection. Individually, only 21-F2 (bin 4) showed any neutralization. All cocktails produced synergistic effects, boosting potencies over an order of magnitude compared with their individual components. All cocktails were mixed in a 1:1:1:1 (4-Ab) or 1:1:1:1:1 (5-Ab) concentration ratio. The 4-membered cocktails contained 23-H7 + 22-D9 (or 21-F2) + 22-E7 (or 22-F7) + 2-A6, while the 5-membered cocktail contained 23-H7 + 22-D9 + 21-F2 + 22-F7 + 2-A6. Thus, all cocktails contained both 23-H7 (bin 2) and 2-A6 (bin S2). The remaining bin 4 and bin C members were: cocktail #1 (4-Ab: TATX-03a) 22-D9 + 22-E7; cocktail #2 (4-Ab: TATX-03b) 21-F2 + 22-F7; cocktail #3 (4-Ab) 22-D9 + 22-F7; cocktail #4 (4-Ab) 21-F2 + 22-E7; and cocktail #5 (5-Ab: TATX-03c) 22-D9 + 21-F2 + 22-F7. Note that cocktail #1 (TATX-03a) and cocktail #3 were each comprised of non-neutralizing Abs (lacked 21-F2). Benchmark Abs used in these assays gave IC50 (µg/ml) values of 0.45 (REGN10987), 0.67 (REGN10933) and 0.58 (REGN10987+REGN10933) in pseudovirus assays and 10.5 (REGN10987), (REGN10933) and 0.3, 0.2 n = 2 (REGN10987+REGN10933) in authentic virus assays.

### Multi-Ab cocktail TATX-03 reduces viral titer in hamster challenge model

To determine the *in vivo* efficacy of TATX-03, two blends and their individual constituent Abs were tested in a Syrian hamster model of acute SARS-CoV-2 infection^15, 16^. Blends TATX-03a and TATX-03b were tested in separate studies, with all Abs (or PBS mock) administered as a single intraperitoneal (i.p.) dose either 24-hours pre-challenge (prophylaxis, PPx) or 4 hours post-challenge (therapeutic, Tx) with SARS-CoV-2 (D614G Mutant BetaCoV/Munich/BavPat1/2020) **(Fig 4a).** Infection was confirmed by PCR on day 1 post-challenge throat swab samples and average daily weight loss was similar for all groups for the duration of the study **(Supplemental Fig S5a-b, Study 2 data not shown).** All animals survived to endpoint at day 4 post-challenge.

**Fig 4.**
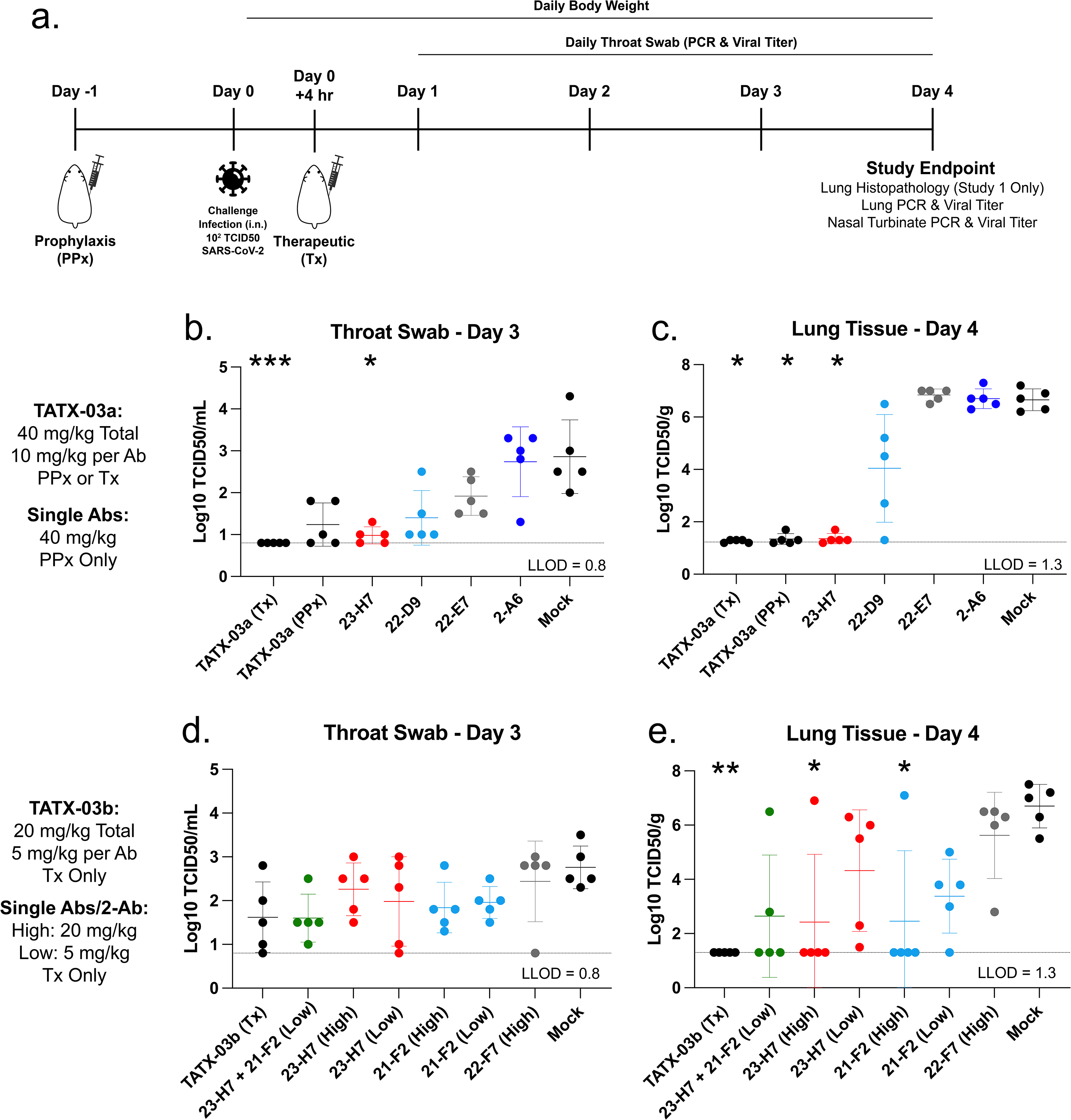
Hamster challenge model for SARS-CoV-2 infection. **(a)** Study design **(see Methods)**. Primary measures of *in vivo* efficacy of cocktails and individual Abs in blends TATX-03a **(b,c)** and TATX-03b **(d,e)**. TATX-03a (23-H7, 22-D9, 22-E7, 2-A6) was administered pre-challenge (prophylactic, PPx) or post-challenge (therapeutic, Tx) as indicated at 40 mg/kg total Ab concentration (10 mg/kg/Ab in the 4-Ab blend), while the individual Abs of TATX-03a were on administered pre-challenge at 40 mg/kg. TATX-03b (23-H7, 21-F2, 22-F7, 2-A6) was administered at 20 mg/kg total Ab concentration (5 mg/kg/Ab in the 4-Ab blend) as Tx only. Individual antibodies were administered post-challenge at 20 mg/kg (23-H7, 21-F2 and 22-F7) and 5 mg/kg (23-H7 and 21-F2). In addition, one group was post-challenge treated with a 2-Ab combination of 23-H7 and 21-F2 at 5 mg/kg total Ab concentration (2.5 mg/kg/Ab). Replication-competent viral titers (Log10 Median Tissue Culture Infective Dose (TCID50) in **(b,d)** throat swab (day 3 post-infection; lowest level of detection, LLOD = 0.8) and **(c,e)** lung (day 4 post-infection, endpoint; LLOD = 1.3). Statistics are described in Methods. *p<0.05, **p<0.01, ***p<0.001.

The first study tested the TATX-03a blend (23-H7, 22-D9, 22-E7, 2-A6), composed of Abs that were non-neutralizing individually but neutralized synergistically as a cocktail in authentic virus *in vitro* **(Fig 3c)**. All animals (5/5) receiving therapeutic administration of the TATX-03a cocktail (40 mg/kg which equals 10 mg/kg/Ab) showed day 3 throat swab virus titers at or below the lowest limit of detection (LLOD), with animals treated with TATX-03a prophylaxis showing clear reductions in day 3 throat swab virus titer compared to mock **(Fig 4b).** At the day 4 endpoint both TATX-03 treated groups demonstrated significantly reduced virus titers in whole lung tissue compared to mock **(Fig 4c)**, with 100% (5/5) of the animals in the therapeutic group and 80% (4/5) of the prophylactically treated animals showing viral titers below LLOD. In line with previous reports of discordant reductions in viral load between lung and nasal turbinate in this model of infection^17^, all but one animal across both studies harbored detectable viral titers in the nasal turbinate at the day 4 endpoint **(Supplemental Fig S5c-d)**, which is anticipated to be due to prominent local infections following intranasal inoculation with a significant bolus of virus. Table 1 summarizes the various replication-competent viral titers per cohort.

**Table 1.** Efficacy results for two different 4-Ab cocktails, TATX-03a (23-H7,22-D9,22-E7,2-A6) and TATX-03b (23-H7,21-F2,22-F7,2-A6) in independent studies (#1 and 2). Antibodies were administered as a single i.p. injection 24-hours pre-challenge (prophylaxis, PPx) or 4-hours post-challenge (therapeutic setting, Tx) at the specified dose (representing total Ab concentration). Five animals were used per cohort. Values are reported for the replication-competent viral titers measured in throat swab at day 3, lung tissue day 4 (end point) and nasal turbinate day 4 (end point). Values represent the mean (± standard deviation) for n = 5. The number of animals per cohort with titers below the lowest limit of detection (LLOD) is also reported as percent and fraction.

| Study # | Ab treatment | PPx or Tx | Dose (mg/kg) | Replication-competent viral titer (Log<sub>10</sub> TCID<sub>50</sub>) |  |  |
| --- | --- | --- | --- | --- | --- | --- |
|  |  |  |  | /ml Throat swab day3 | /g Lung tissue day4, end point | /g Nasal turbinate day4, end point |
| 1 | 4-Ab (TATX-03a) | Tx | 40 | 0.8 (0), 100% (5/5) LLOD | 1.26 (0.05), 100% (5/5) LLOD | 6.72 (0.44) |
| 1 | 4-Ab (TATX-03a) | PPx | 40 | 1.24 (0.52), 40% (2/5) LLOD | 1.34 (0.21), 80% (4/5) LLOD | 5.8 (0.45) |
| 1 | 3-Ab (23-H7,22-D9,22-E7) | PPx | 40 | 0.98 (0.30), 60% (3/5) LLOD | 3.18 (2.58), 60% (3/5) LLOD | 5.78 (1.51) |
| 1 | 2-Ab (23-H7,22-D9) | PPx | 40 | 1.08 (0.31), 40% (2/5) LLOD | 2.28 (2.30), 80% (4/5) LLOD | 5.14 (1.52) |
| 1 | 23-H7 | PPx | 40 | 0.98 (0.20), 40% (2/5) LLOD | 1.36 (0.19), 80% (4/5) LLOD | 5.7 (0.41) |
| 1 | 22-D9 | PPx | 40 | 1.4 (0.65) | 4.04 (2.06), 20% (1/5) LLOD | 7.2 (0.31) |
| 1 | 22-E7 | PPx | 40 | 1.92 (0.46) | 6.84 (0.23) | 7.8 (0.61) |
| 1 | 2-A6 | PPx | 40 | 2.74 (0.83) | 6.7 (0.37) | 7.7 (0.47) |
| 1 | Mock (PBS) | PPx | 0 | 2.86 (0.88) | 6.66 (0.42) | 6.48 (2.29), 20% (1/5) LLOD |
| 2 | 4-Ab (TATX-03b) | Tx | 20 | 1.62 (0.81) | 1.3 (0), 100% (5/5) LLOD | 6.14 (1.27) |
| 2 | 4-Ab (TATX-03b) | Tx | 5 | 2.32 (0.66) | 4.22 (2.71) | 7.48 (0.64) |
| 2 | 2-Ab (23-H7,21-F2) | Tx | 5 | 1.6 (0.55) | 2.64 (2.25), 60% (3/5) LLOD | 6.14 (1.80) |
| 2 | 23-H7 | Tx | 20 | 2.26 (0.60) | 2.42 (2.50), 80% (4/5) LLOD | 6.5 (1.61) |
| 2 | 23-H7 | Tx | 5 | 1.98 (1.02) | 4.32 (2.25) | 7.32 (0.79) |
| 2 | 21-F2 | Tx | 20 | 1.84 (0.58) | 2.46 (2.59), 80% (4/5) LLOD | 6.36 (1.84) |
| 2 | 21-F2 | Tx | 5 | 1.96 (0.36) | 3.38 (1.36), 20% (1/5) LLOD | 6.24 (1.90) |
| 2 | 22-F7 | Tx | 20 | 2.44, 20% (1/5) LLOD | 5.62 | 7.86 |
| 2 | Mock (PBS) | Tx | 0 | 2.76 | 6.70 | 8.26 |
|  |  |  |  | LLOD = 0.8 | LLOD = 1.3 | LLOD = 2.4 |

Histopathological analysis of airway tissues harvested at the day 4 endpoint showed relatively minimal changes in gross pathology (as expected with the study endpoint coinciding with the acute phase of disease), however, the severity/extent of immune cell infiltration was reduced in groups treated with the TATX-03a cocktail compared to mock, resulting in reduced bronchitis and tracheitis severity scores **(Fig 5c-d).** Fig 5a and b show representative images of H&E-stained lung tissue with bronchitis severity score 0 and 3, respectively.

**Fig 5.**
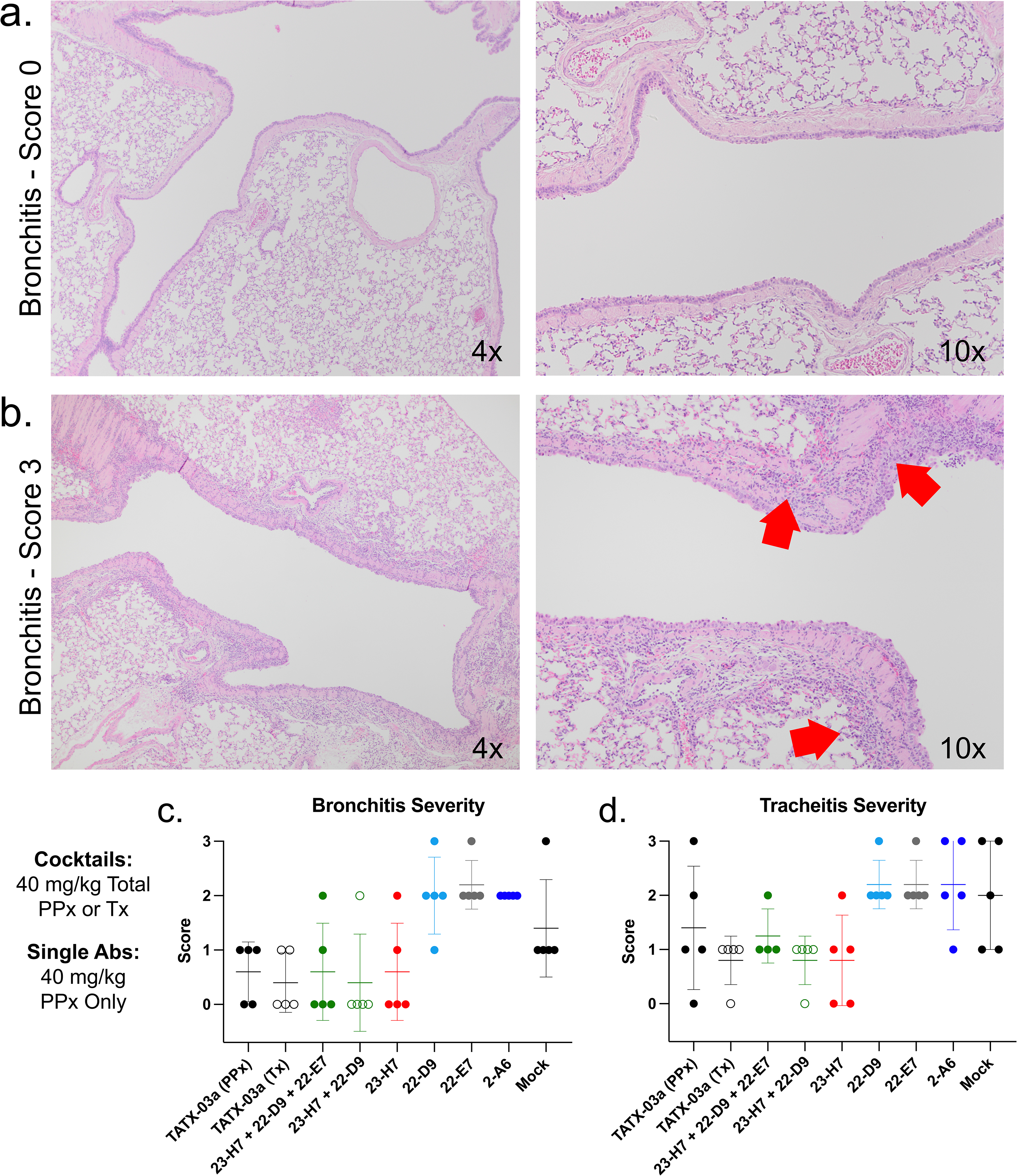
Histopathology analysis of challenge infection model. **(a,b)** Representative images from hematoxylin and eosin (H&E) stained slides of endpoint lung tissue shown at multiple magnifications for bronchitis scores of 0 **(a)** and 3 **(b).** Red arrows point at areas of significant inflammatory cell infiltration. **(c,d)** Average bronchitis **(c)** and tracheitis **(d)** severity as scored by an independent pathological assessment. Scores were determined by the extent of inflammatory cell infiltration into the tissue section.

Two individual Abs demonstrated partial (22-D9) or significant (23-H7) efficacy by these measures despite their inability to neutralize authentic virus *in vitro*; notably, 80% (4/5) of the animals prophylactically treated with 23-H7 achieved undetectable viral load in the lung (below LLOD), with the fifth animal showing viral titer barely above LLOD.

The second blend tested in *vivo*, TATX-03b (23-H7, 21-F2, 22-F7, 2-A6), included the only Ab that neutralized authentic virus *in vitro* **(Fig 3c).** To determine whether *in vitro* synergy could be recapitulated *in vivo*, Abs were administered individually either at a high dose (20 mg/kg) or at a low dose (5 mg/kg), with the latter matching their individual contributions to the TATX-03b cocktail (dosed at 20 mg/kg total Ab, equivalent to 5 mg/kg/Ab). Like TATX-03a, all animals (5/5) treated with the TATX-03b showed undetectable viral titers in lung at the day 4 endpoint. Only high dose monotherapy with 23-H7 or 21-F2 resulted in viral titer reduction in lung comparable to the cocktail in 80% (4/5) of the animals, while viral titer reduction was clearly less pronounced in animals treated with the corresponding low dose monotherapy **(Fig 4e, Table 1)**. This indicates that the cocktail’s efficacy cannot not solely be attributed to the presence of 23-H7 or 21-F2 individually, strongly suggesting a synergistic effect. When dosed as a 2-Ab cocktail (23-H7, 21-F2) at 5 mg/kg total Ab concentration (equivalent to 2.5 mg/kg/Ab), viral load in the lung at the day 4 endpoint was reduced to undetectable levels in 60% (3/5) of animals, which was more efficacious than either individual Ab when administered at 5 mg/kg, consistent with a synergistic effect **(Fig 4e).**

### Cocktail formulation overcomes escape of individual Abs by variants of concern (VOCs)

To determine whether the components of the TATX-03 cocktail were resistant to SARS-CoV-2 VOCs, individual Abs were screened against a panel of cell-associated spike trimers harboring mutations from the B.1.1.7 (Alpha), B.1.351 (Beta), P.1 (Gamma), B.1.429, and B.1.526 lineages, as well as the A (Wuhan-1) and B (D614G) parental lineages for reference. These data revealed a bin-dependent susceptibility to viral variants **(Fig 6a)**, which generally was in line with the data obtained from ELISA-based reactivity screening towards plate-adsorbed recombinant spike variants **(Supplement Fig S6a-c).** Bin 2 (23-H7) was resistant to all cell-associated spike trimer variants tested except for those carrying the L452R mutation (B.1.429), which results in reduced, but not totally abrogated, binding of Bin 2. No other Abs in the cocktail were susceptible to B.1.429. Bin 4 Abs (22-D9 and 21-F2) were similarly susceptible to B.1.351 but differentially susceptible to B.1.526. Also, while both 22-D9 and 21-F2 showed no binding to the P.1 variant in the context of cell-associated spike, 22-D9 retained low-level binding to P.1 by ELISA. Bin C (22-E7 and 22-F7) was uniformly susceptible to B.1.1.7 and B.1.526 with no difference observed towards other mutants. Bin S2 (2-A6) showed uniform binding to all tested mutant trimer constructs, being unaffected by any of the mutations in the S2 subunit. Bin C (22-E7 and 22-F7) and bin S2 (2-A6) clones both showed some reactivity discrepancies towards cell-associated spike compared to their binding profiles against plate-immobilized soluble recombinant spike as revealed by ELISA, which is likely due to the trimer being cleavable on cells, representing a more “native” conformation, contrasting the more artificial cleavage-resistant stabilized forms of the soluble recombinant constructs. Overall, no two bins represented in the TATX-03 cocktail could be simultaneously escaped by the variants tested.

**Fig 6.**
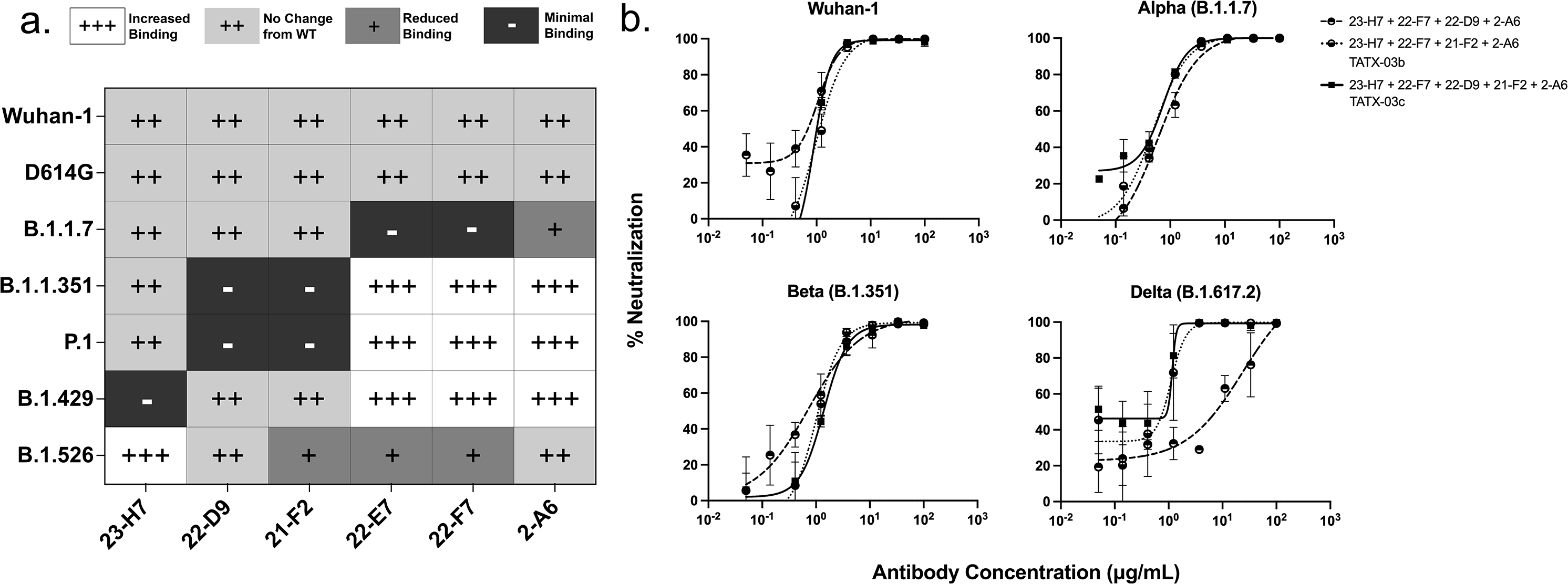
Cell-based reactivity and pseudovirus neutralization screening against the variants of concern (VOCs) **(a)** Heat map summarizing the complementary vulnerabilities of indicated clones of the TATX-03 cocktail against cell-associated spike trimers expressing Alpha (B.1.1.7), Beta (B.1.351), Gamma (P.1), B.1.429, and B.1.526 mutants. Corresponding dose-response graphs are shown in **Supplemental Fig 6a**. **(b**) *In vitro* virus neutralization screenings of TATX-03a-c c using VSV-particles pseudo-typed with spike proteins representing the original Wuhan-1 isolate and SARS-CoV-2 variants of concern Alpha, Beta, and Delta..

To assess the functional consequence of mutant susceptibility, we tested whether TATX-03 could retain neutralization potency in pseudovirus assays adapted to the Alpha, Beta, and B.1.617.2 Delta variants. In addition to the two TATX-03 4-Ab cocktails, a 5-Ab version of TATX-03 (TATX-03c) was tested which included two bin 4 antibodies (21-F2 and 22-D9) to potentially capitalize on subtle differences in their binding specificities identified by the epitope binning experiments and VOCs screenings. All analyzed multi-Ab cocktails retained potent neutralization against the tested pseudotyped viruses **(Fig 6b)** with only one 4-Ab combination showing partial susceptibility to the Delta variant.

## DISCUSSION

Globally to date (as of July 2021), there have been over 189 million COVID-19 cases and over four million related deaths, of which over 0.6 million deaths have occurred in the United States (Johns Hopkins Center for Systems Science and Engineering). While mass vaccination is underway, more effective and sustainable treatment strategies are needed to prevent and treat COVID-19. Antibodies may provide significant benefit in treating newly SARS-CoV-2-infected individuals with high-risk for progression to severe disease. To date, four antibody therapeutics targeting the SARS-CoV-2 spike protein have received emergency use authorization (two monoclonal antibodies, and two two-Ab combination therapies). However, the exquisite specificity of Abs also renders them susceptible to mutagenic escape. Combining Abs into multi-Ab cocktails in order to broaden the aggregate epitope footprint by targeting diversified epitopes is anticipated to significantly reduce this mutagenic escape risk. Here, we have demonstrated the use of pre-existing fully human B cell libraries to rapidly identify a highly diverse panel of SARS-CoV-2 spike protein-specific Abs for curation into therapeutic cocktails capable of resisting mutagenic escape.

Intriguingly, the pair of RBD-specific binders used in our TATX-03 cocktail (bin 2 + bin 4) target similar epitopes as those incorporated in dual therapies with reported *in vivo* efficacy that have been discovered independently by other groups; REGN10987 (overlaps our bin 2) + REGN10933 (overlaps our bin 4); CoV2-06 + CoV2-14; and COV2-2130 + COV2-2196^6;18–21^. While this suggests that only particular combinations of Abs will produce *in vivo* protection, these therapies are more susceptible to mutagenic escape and may only utilize neutralization as the primary mechanism of action. To our knowledge, we are the first to report on the discovery of a multi-Ab cocktail, beyond a dual therapy, that produces *in vivo* protection against SARS-CoV-2 via synergistic effects in an animal model of infection.

Authentic virus neutralization assays are often used as a triage step for prioritizing Abs for *in vivo* studies, however, by this criteria, 23-H7 would have been overlooked. Since we intentionally retained high functional diversity in our leads, we progressed this clone due to its uniquely high affinity, kinetic perturbation/partial blockade of ACE2 binding, potency in pseudovirus assay and ability to pair with other neutralizing and non-neutralizing bins in cocktails. This resulted in its prioritization for *in vivo* testing where it was highly efficacious as both a monotherapy and in combination with other Abs in prophylactic or therapeutic settings. While the discrepancy between our *in vitro* pseudovirus- and authentic virus-based neutralization screening data might indicate a bias towards strong ACE2 blocking potency for the latter assay format, the lack of correlation between *in vitro* neutralization and *in vivo* protection has been reported by others^7^, and suggests that non-neutralizing mechanisms of action can provide therapeutic benefit. We anticipate that the high human IgG1 decoration degree of the spike trimer, resulting from simultaneous binding of the individual cocktail components, contributes to *in vivo* viral load reduction by enhancing Fc-mediated clearance.

The breadth of our discovery approach not only resulted in opportunities for combining multiple Abs across distinct non-overlapping bins, but also enabled us to leverage nuanced intra-bin variations, facilitating rapid reformulation of cocktails in response to mutations or seasonal variations in the SARS-CoV-2 genome. Indeed, our detailed binning analysis suggested that 22-D9 and 21-F2 bind different, but overlapping epitopes, which was corroborated by their differential susceptibility to cell-associated spike of the B.1.526 lineage, so we retained both in a 5-Ab cocktail formulation to provide a broader resistance to viral variants, while retaining potent neutralization via synergistic effects. A similar compensation strategy was likely the rationale for combining bamlanivimab and etesevimab, which bind different but overlapping epitopes on RBD^9^. However, applying this strategy to a dual therapy, resulting in a “refined” monotherapy that is still vulnerable to escape by a single point mutation.

In conclusion, we are the first to describe a multi-Ab cocktail, beyond dual therapy, that shows synergistic effects both *in vitro* and *in vivo* and confers protection in an *in vivo* animal model of SARS-CoV-2 infection in both prophylactic and therapeutic settings. Our findings support the use of both neutralizing and non-neutralizing mechanisms of action in our synergistic TATX-03 multi-Ab cocktail to retain efficacy towards an evolving viral genome, meriting its investigation in the clinic as a potential therapeutic intervention for SARS-CoV-2 infection.

## METHODS

### Recombinant proteins

Various targets were used as panning, screening and analytical reagents for ELISA and Octet binding assays and purchased from different vendors. SARS-CoV-2 spike trimer, RBD-hFc, RBD (tagless), human ACE2-hFc, and SARS-CoV-1 spike trimer were produced using the rPEx^TM^ recombinant protein production platform. In brief, HEK293-EBNA1 cells were transiently transfected with pUPE expression vectors containing recombinant protein encoding, codon-optimized synthetic genes. Six days post transfection secreted recombinant proteins were purified to homogeneity by a two-step purification approach, including affinity purification and Size-Exclusion Chromatography. SARS-CoV-2 spike trimer (cat# A33-11-02-SMT1) was obtained from the National Research Council (Quebec, Canada). SARS CoV-2 full-length spike protein and B.1.1.7, B.1.351, P.1, B.1.429, B.1.617 and B.1.525 mutations were purchased form Cube Biotech (cat# 28702, 28717, 28720, 28723, 28736, 28739, and 28733). Various spike protein subunits including S1-mFc, S1-hFc, S2-hFc, NTD-hFc, S1-S2-His, S1-His, and mutants S1-His(D614G), S. African B.1.351 lineage S1-His(K417N, E484K, N501Y, D614G), S1-His(HV69-70del, N501Y, D614G), UK S1-His(HV69-70del, Y144del, N501Y, A570D, D614G, P681H) were purchased from Sino Biological (cat# 40591-V05H1, 40591-V02H, 40590-V02H, 40591-V41H, V0589-V08B1, 40591-VO8H, 40591-V08H3, 40591-V08H10, 40591-V08H7, 40591-V08H12, respectively) as well as RBD single point mutants (A435S, F342L, G476S, K458R, N354D, N439K, S477N, V367F, V483A, W436R, E484K, K417N, Y453F, N501Y; cat# 40592-V08H4, 40592-V08H6, 40592-V08H8, 40592-V08H7, 40592-V08H2, 40592-V08H14, 40592-V08H46, 40592-V08H1, 40592-V08H5, 40592-V08H9, 40592-V08H84, 40592-V08H59, 40592-V08H80, 40592-V08H82, respectively).

Benchmark Abs were internally generated from published sequences and expressed in the human IgG1 framework resembling the one used for the Ab candidates: B38^5^ (PDB ID: 7BZ5), CB6^4^, CR3022^23^ H4^5^, REGN10933 & REGN10987^6^, or as VHH-hFc1: VHH-72^24^, SB14^25^, and SB68^26^. Bococuzimab (hIgG1) was produced as isotype control antibodies. HRP-conjugated anti-hKappa and anti-hLambda detection reagents were purchased from SouthernBiotech. Antibodies, expressed either as human IgG1-Fab and full-length human IgG1, destined for (initial) *in vitro* binding experiments were produced following transient transfection of HEK293 cells. while full-length human IgG1 antibodies designated for *in vitro* neutralization and animal studies were produced in transiently transfected and/or stable CHO cells. Fabs were purified by a CH1 matrix resin, while IgGs were purified by protein A chromatography. Antibodies destined for animal studies were produced in CHO cells. All Fabs and IgG1s were subjected to an additional purification step in-house using gel filtration and formulated in PBS.

### Library panning

The phage display libraries employed here were previously generated by ImmunoPrecise Antibodies (Naïve Human Library #0899, Autoimmune Patient Library #0845, and Llama VHH Library #3566). Briefly, B lymphocytes were isolated from the indicated biological source, and Ab variable region sequences were amplified by RT-PCR. The fragments were ligated into the pHENIX-His8-VSV vector and transformed into *Escherichia coli* TG1 cells. (Sub)Library rescue was conducted prior to each round of antigen panning by inoculating bacterial cells into TYAG medium followed by the addition of helper phage to induce phage production. Phage particles were isolated by PEG/NaCl precipitation and filtered using a 0.45 µm filter.

Either magnetic Streptavidin, Protein A or polystyrene beads were coated with the (biotinylated) antigen of interest and washed to remove any unbound protein. Purified phage particles were blocked in PBS + 5% skim milk and any bead-reactive Abs were depleted by pre-incubation with uncoated beads. Unbound phages were subsequently incubated with the antigen-coated beads, followed by washing in PBS-Tween to remove any unbound phage particles. Depending on the panning strategy, either the bead-bound (for positive antigen selection) or unbound (for negative selection) phage particles were incubated with TG1 cells followed by phage rescue by helper phage superinfection. For target enrichment, eleven unique panning strategies were conducted in parallel using varying combinations of S1, S2, and RBD subunits of the SARS-CoV-2 spike protein, and a fully assembled, stabilized spike trimer of the related SARS-CoV-1). Depletion panning with human ACE2- or CR3022-bound spike (where CR3022 represents an anti-SARS-CoV-1 antibody from the literature with cross-reactivity to SARS-CoV-2^23, 27^) and irrelevant non-target proteins were performed to steer towards binders to specific epitopes and increase target specificity by reducing off-target reactivity, respectively.

### ELISA screening

Targets diluted to final concentrations 1.5 µg/ml in carbonate binding buffer were added to Greiner Bio-One High Bind ELISA plates in 50 µL/well and incubated overnight at 4°C. Plates were blocked with 1% BSA in PBS for 60 minutes. If an additional capture step was performed it was conducted in PBS for 1 hour at room temperature (RT). Coated plates were finally washed with PBS-T before serial dilutions of phage or recombinant Abs were added in duplicate in PBS supplemented with 5% skim milk or 1% BSA and incubated at RT for 60 minutes. After washing with PBS-T, appropriate HRP-conjugated secondary antibody (anti-M13 and goat-anti-human-IgG for detection of phages and full size hIgG1, repsectively) was added and incubated for 60 minutes at RT. For the initial reactivity screenings of scFv-containing periplasmic fraction, target-bound scFvs were detected by incubation with mouse anti-VSV followed by anti-mouse IgG-HRP. After final washing wells were typically stained with 50 µL TMB substrate for 10 minutes and the reaction was stopped by adding 50 µL 2M H2SO4. Absorbance was read at 450 nm on an Envision multimode plate reader and data was processed in GraphPad Prism.

### Interaction analysis by Octet

All label-free interaction analyses were performed on an Octet HTX biolayer interferometry-based detection system (ForteBio/Sartorius, Gottingen, Germany) equipped with various sensor types; AR (amine-reactive), SAX (streptavidin-coated), or AHC (anti-human-Fc capture) sensors. Experiments were conducted at 25°C in a run buffer of PBS + 0.05% Tween-20 + 0.1% BSA.

#### Binding affinity estimates

Different assay formats were used to estimate the binding affinities of Ab/target bimolecular interactions. In one assay format, Fab was titrated as monovalent analyte (typically as a 3-fold dilution series with a top concentration of 3 µM, and at least one concentration in duplicate) over AHC sensors coated with human-Fc-fused targets RBD-hFc, S1-hFc or S2-hFc as ligands (Sino Biological). In the reverse format, tagless RBD or S1-His(D614G) were titrated as monovalent analytes over Ab-coated AHC sensors. Global affinity estimates were determined using the Kinetics module of Fortebio’s Data Analysis HT software version 12.0.1.55. Data were processed by subtracting the responses of a buffer analyte sample and fitting these referenced data globally to a simple 1:1 Langmuir binding model to deduce the K_D_ value from the ratio of the kinetic rate constants (K_D_ = k_d_/k_a_), where k_d_ and k_a_ are the dissociation and association rate constants, respectively. Interactions showing square-shaped binding curves were alternatively fit to a steady-state (equilibrium) isotherm; affinities deduced from kinetic and equilibrium fitting routines were equivalent.

Additionally, the solution affinity of the 23-H7-Fab/RBD binding interaction was determined by titrating 23-H7 Fab (1000-1.4 nM, as a 7-membered 3-fold dilution series) into RBD fixed at 5 nM, allowing these solutions to equilibrate (an hour at RT) and then probing for free RBD in these samples using SAX sensors coated with biotinylated-23-H7-IgG. All samples were measured on duplicate sensors. An apparent solution affinity (or IC50 value) was determined by fitting the reference-subtracted responses (from a buffer analyte sample) to a non-linear regression, inhibition dose-response curve (4-parameter least-squares fit) model in GraphPad Prism software version 9.

#### Pairwise epitope binning

Combinatorial pairwise Ab competition or “epitope binning” assays were performed using various assay formats. To perform a “classical sandwich” assay format, Abs were covalently coupled onto AR sensors using standard coupling conditions to generate the ligands (surface-immobilized Abs) and used to capture S1-His(D614G) monovalent target (typically 5 µg/ml, 65 nM) followed by an Ab analyte typically at 10 µg/ml (133 nM binding sites). Alternatively, reaction surfaces were generated by coating SAX sensors with 5 µg/ml biotinylated Abs. Ligands were regenerated with 75 mM phosphoric acid. “Waterfall” experiments were conducted on freshly Ab-coated SAX sensors (single use, not regenerated) using 5 µg/ml S1-His(D614G) followed by an Ab titration spanning 6000 –25 nM binding sites as a 6-membered 3-fold dilution series, with one concentration (667 nM) in duplicate. Data were analyzed in the Epitope Binning molecule of Fortebio’s Data Analysis HT software version 12.0.1.55. Heat maps were curated manually in Excel by merging the results from different experiments.

#### Multi-Ab epitope binning

To perform a “tandem cocktail” multi-Ab binning experiment, SAX sensors were coated with 5 µg/ml biotinylated 23-H7 (bin 2) and used to tether 5 µg/ml spike trimer. Three Ab analytes from non-overlapping bins were associated in consecutive analyte binding steps, each step building upon the complex formed in the previous steps. For example, bin 4 Ab was used in step 1, bin 4+C was used in step 2, and bin 4+C+S2 was used in step 3, thereby maintaining saturating levels of the Ab analyte applied in the previously applied steps to eventually saturate the 23-H7-tethered spike with three Ab analytes (from bins 4, C and S2). The responses of each newly applied Ab analyte to “Ab-saturated” 23-H7-tethered spike were compared with the responses of that Ab analyte to the “naked” 23-H7-tethered spike. Data were processed in ForteBio’s Data Acquisition software version 12.0.1.8 by Y-aligning to zero at each association step.

Alternatively, multi-Ab binnings were performed in a “premix” assay format. To prepare the reaction surfaces for these experiments, SAX sensors were coated with 5 µg/ml biotinylated Abs from different bins (e.g., 2, 4, C, or S2) or with controls; biotinylated ACE2-hFc or mouse anti-His mAb (R&D systems). Spike trimer (1 µM binding sites) was premixed with Abs from different epitope bins, either individually, or as 2-, 3-, or 4-membered cocktails using Abs at saturating concentrations (10 µM binding sites). Samples of premixed spike/Ab complexes, spike alone or buffer were used as analytes for binding to the Ab-coated sensors (or control surfaces) to probe for free binding sites in these mixtures. Binding responses were compared with those of spike alone and determined to be blocked if their responses were significantly suppressed to baseline levels (like the buffer blank).

### Cell-associated spike screening

Synthetic genes encoding for SARS-CoV-2 surface glycoprotein variants, including B.1.1.7 (Alpha), B.1.351 (Beta), P.1 (Gamma), B.1.429, B.1.526 and B.1.617 (Delta) lineages, as well as the A (Wuhan-1) and B (D614G) parental lineages, were obtained from GeneArt and subsequently cloned into IPA’s mammalian expression vector. To induce expression of spike trimers in a cell context, HEK293F cells were transiently transfected using FectoPRO per the manufacturer specification (PolyPlus Transfection, Illkirch, France) with SARS-CoV-2 surface glycoprotein expression vector. 48 hours post transfection, cells were harvested, washed and dispensed to 96-well cell culture plates at a concentration of 1.0×10^5^ cells per well, and serial dilutions of test or control Abs were added in a final volume of 30 µL per well in duplicate. After 1 hour the wells were washed and Ab binding was detected with Donkey F(ab’)2 Anti-Human IgG conjugated to phycoerythrin (Abcam cat.# ab102439). Cells were analyzed using an iQue High-Throughput Flow Cytometer (Sartorius, Göttingen, Germany). EC50 values were calculated in GraphPad Prism.

### Pseudovirus neutralization

The production of VSV virus particles expressing the SARS-CoV-2 Spike protein has been previously described^28^. Briefly, the SARS-CoV-2 Spike protein was cloned into the pCAGGS expression vector system and transfected into HEK-293T cells. Cells were then infected with the VSVΔG pseudotyped virus further modified to encode the *Photinus pyralis* luciferase reporter protein. After 24 hours supernatants were collected and titrated on African green monkey VeroE6 cells. In neutralization assays, Abs were diluted in DMEM supplemented with 1% fetal calf serum (Bodinco), 100 U/ml Penicillin, and 100 µg/ml streptomycin before being added to an equal volume of pseudotyped virus particles and incubated at RT for 1 hour. The mixture was then added to a confluent monolayer of VeroE6 cells in a 96-well tissue culture plate and incubated for 24 hours. Following this incubation luciferase activity was measured in the presence of D-luciferin substrate (Promega) using a Centro LB960 plate luminometer (Berthold). Neutralization was calculated as the ratio of luciferase activity in the presence of Abs normalized to a negative control well containing only pseudotyped virus and no Abs.

### Authentic virus neutralization

Authentic virus neutralization assays were performed at ViroClinics Biosciences (Rotterdam, Zuid-Holland) using the SARS-CoV-2 virus (BetaCoV/Munich/BavPat1/2020) carrying the D614G mutation. In short, per sample a two-fold serial dilution was incubated with a fixed amount of virus (200 TCID_50_/well or 4000 TCID_50_/mL) in triplicate for 1 hour at 37 °C with a starting Ab concentration of 100 µg/mL. Next, the virus-Ab mixtures were transformed to plates with VeroE6 cell culture monolayers and after the incubation period of 5-6 days at 37 °C cytopathic effect (CPE) in the monolayer was measured and scored by the vitality marker WST8 and neutralization titers were calculated according to the Reed-Muench method^29^.

### *In vivo* hamster challenge model of infection

All animal studies were performed at ViroClinics Xplore, Schaijk, The Netherlands and conducted according to European Union Directive 2010/63/EU and the standards of Dutch law for animal experimentation. Ethical approval for the studies is registered under project license number AVD277002015283-WP26 and -WP33. The animal model studies were conducted as previously described^30^. Briefly, groups of five specific pathogen free (SPF) male Syrian golden hamsters (Mesocricetus auratus) aged 9 to 10 weeks at the start of the experiment were randomly assigned to experimental groups. Animals were housed in elongated Type 2 cages with two or one animal per cage (as five animals per group) under BSL-III conditions during the experiment. Antibody or mock (PBS) treatment were administered as a single intraperitoneal injection at the indicated time. All animals were challenged at day 0 with a single intranasal administration of 10^2.0 TCID50 SARS-CoV-2 (BetaCoV/Munich/BavPat1/2020 kindly provided by Prof. Dr. C. Drosten) in a total dose volume of 100 µL administered equally over both nostrils. On day 4 post-challenge all animals were euthanized by abdominal exsanguination under isoflurane anesthesia (3-5%).

### Animal study tissue collection

Animals were weighed and throat swabs were collected daily post infection. At the time of euthanasia, lung lobes were inspected and observed percentage of affected lung tissue was estimated, samples of the left nasal turbinates, trachea and the entire left lung (often with presence of the primary bronchi) were preserved in 10% formaldehyde for histopathology and samples of the right lung parenchyma and right nasal turbinates were collected. Throat swabs and right lung and nasal turbinate tissues were frozen for subsequent virological assessment by quantitative PCR and virus titration.

### Viral load quantification from *in vivo* samples

For determination of replication competent virus levels, quadruplicate 10-fold serial dilutions were used to determine the virus titers in confluent layers of Vero E6 cells. In short, serial dilutions of the samples (throat swabs and tissue homogenates) were prepared and incubated on Vero E6 monolayers for 1 hour at 37 °C. Vero E6 monolayers are washed and incubated for 5 or 6 days at 37 °C. Viability was measured by scoring using the vitality marker WST8. Viral titers (log10 TCID50/ml or /g) were calculated using the method of Spearman-Karber. For detection of viral RNA levels in the samples, RNA was extracted from samples using Magnapure LC total nucleic acid isolation kit (Roche). RNA amplification and quantification were carried out using a 7500 Real-Time PCR System (Applied biosystems) specific primers (E_Sarbeco_F: ACAGGTACGTTAATAGTTAATAGCGT and E_Sarbeco_R:ATATTGCAGCAGTACGCACACA) and probe (E_Sarbeco_P1: ACACTAGCCATCCTTACTGCGCTTCG) as described previously^31^ and RNA copies (log10 copies/ml or /g) were calculated.

### Histopathological evaluation of tissue from *in vivo* studies

After fixation with 10% neutral-buffered formalin, lung, nasal turbinate and trachea tissues were sectioned, paraffin embedded, micro-sectioned to 3 μm on glass slides and stained with hematoxylin and eosin for histopathological evaluation. The stained tissues were examined by light microscopy using an Olympus BX45 light microscope with magnification steps of 40x, 100x, 200x, and 400x for scoring. Severity of inflammation was scored based on inflammatory cell infiltration in tracheas and bronchi (0 = no inflammatory cells, 1 = few inflammatory cells, 2 = moderate number of inflammatory cells, 3 = many inflammatory cells).

### Statistical Analysis

All treatment groups were compared with the mock group. The treatment groups were compared on the development of weight, throat swab RT-PCR and throat swab virus titration. Mixed model analyses were conducted in SAS with Proc Mixed. A Dunnet correction for multiple testing was applied. For the virology and histopathology variables measured on day 4 post-challenge a two-sided p-value was calculated for Fisher’s Exact Test for categorical variables and the Wilcoxon Rank Sum Exact Test for continuous and ordinal variables. Since the statistical analysis of these variables was explorative in nature, no correction for multiple testing was used. For values below the lower limit of detection the lower limit of detection was used.

## Acknowledgements

The authors are gratefully thankful to all IPA employees who have contributed with passion and unlimited effort to the work that has led to the results published in this paper.

## SUPPLEMENTAL FIGURE LEGENDS

**Fig S1.**
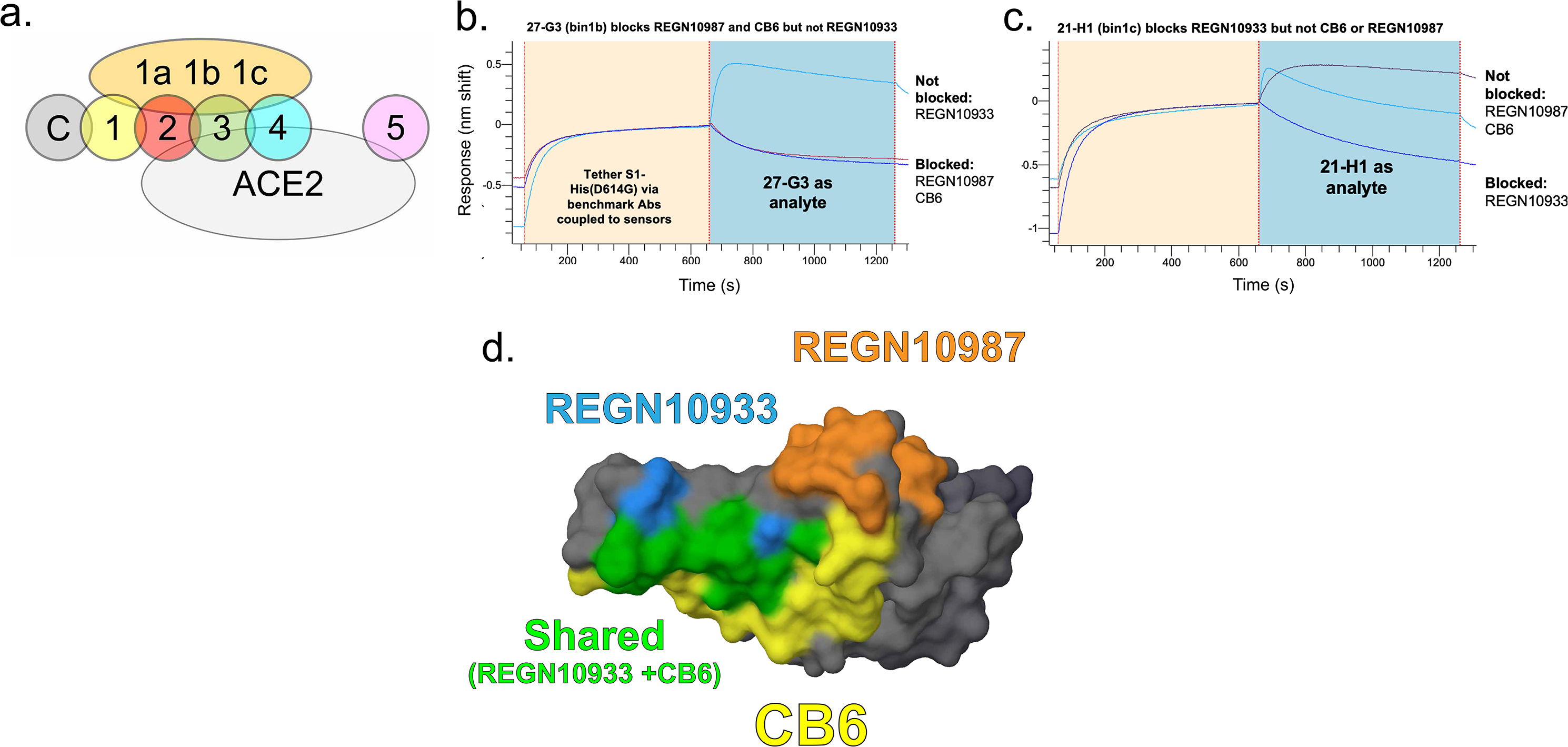
Additional data from pairwise epitope binning assays. (**a-c**) Nuanced binning profiles for the sub-bins including a Venn diagram demonstrating cross-blockade between bins **(a)**. Sensorgram overlay plots showing the sandwich binding of **(b)** 27-G3 (bin 1b) or **(c)** 21-H1 (bin 1c) as analyte to recombinant S1-His(D614G) that is first tethered via sensors coated with benchmark Abs. In this example, 27-G3 (bin 1b) blocks REGN10987 and CB6 but not REGN10933, whereas 21-H1 (bin 1c) blocks REGN10933 but not CB6 of REGN10987. **(c)** Image of the RBD in grey (PDB ID: 7BNN residues 330-520) showing the epitope contacts for benchmark Abs REGN10987 (orange, PDB ID: 6XDG), REGN10933 (blue, PDB ID: 6XDG), CB6 (yellow, PDB ID: 7C01), and REGN10933/CB6 shared residues in green. Despite REGN10933 and CB6 sharing substantial overlapping epitope contacts, our library contained clones that discriminated between them, as here shown by bin 1b and bin 1c.

**Fig S2.**
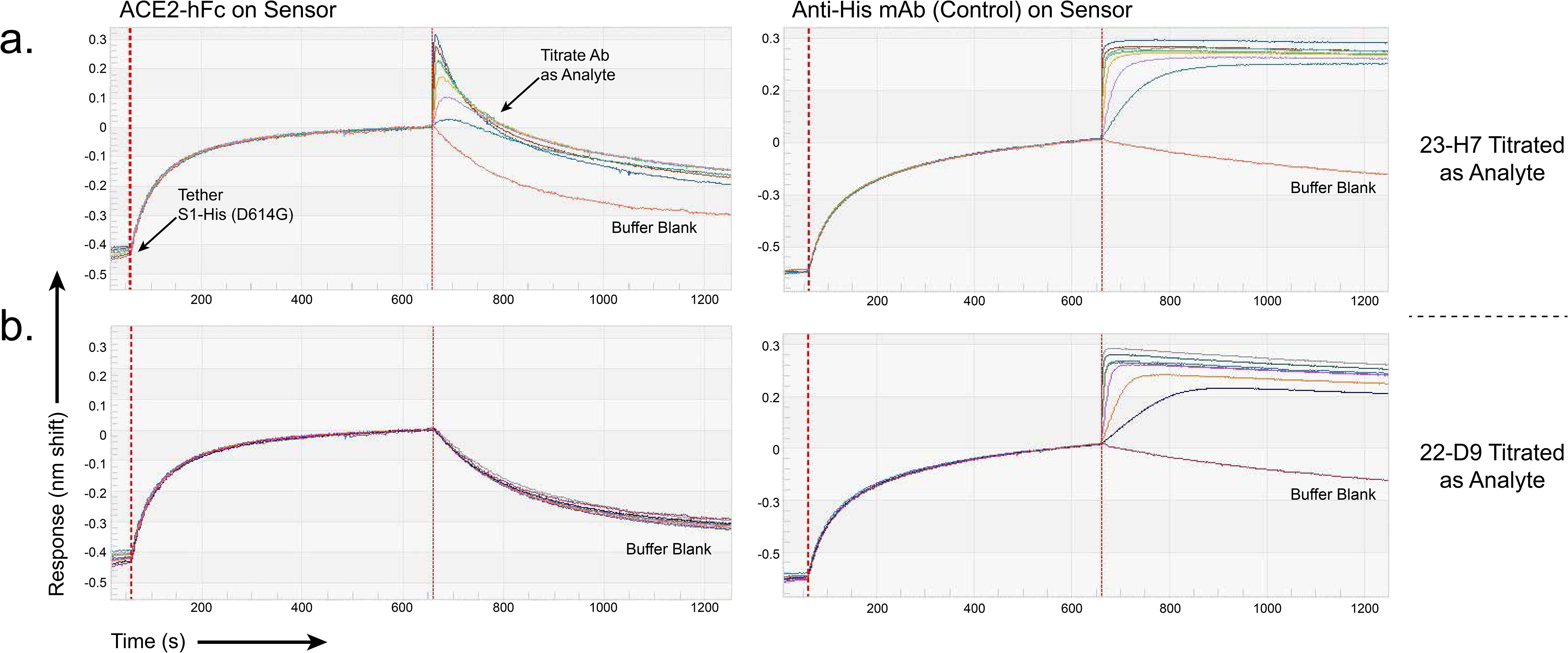
“Waterfall” classical binning assays. Titration of analytes **(a)** 23-H7 or **(b)** 22-D9 over S1-His(D614G) that is tethered via ACE2-coated sensors (left) showing that 23-H7 binding is kinetically perturbed (partially blocked) suggesting that 23-H7 and ACE2 target closely adjacent or minimally overlapping epitopes, whereas 22-D9 is fully blocked, suggesting that its epitope may overlap substantially with that of ACE2. Dose-dependent unhindered sandwiching signals of S1-His(D614G) tethered via anti-His mAb-coated sensors (right) serve as controls to indicate the results expected for analyte binding at a distinctly different and non-overlapping site relative to that of the immobilized binding partner on the sensor.

**Fig S3.**
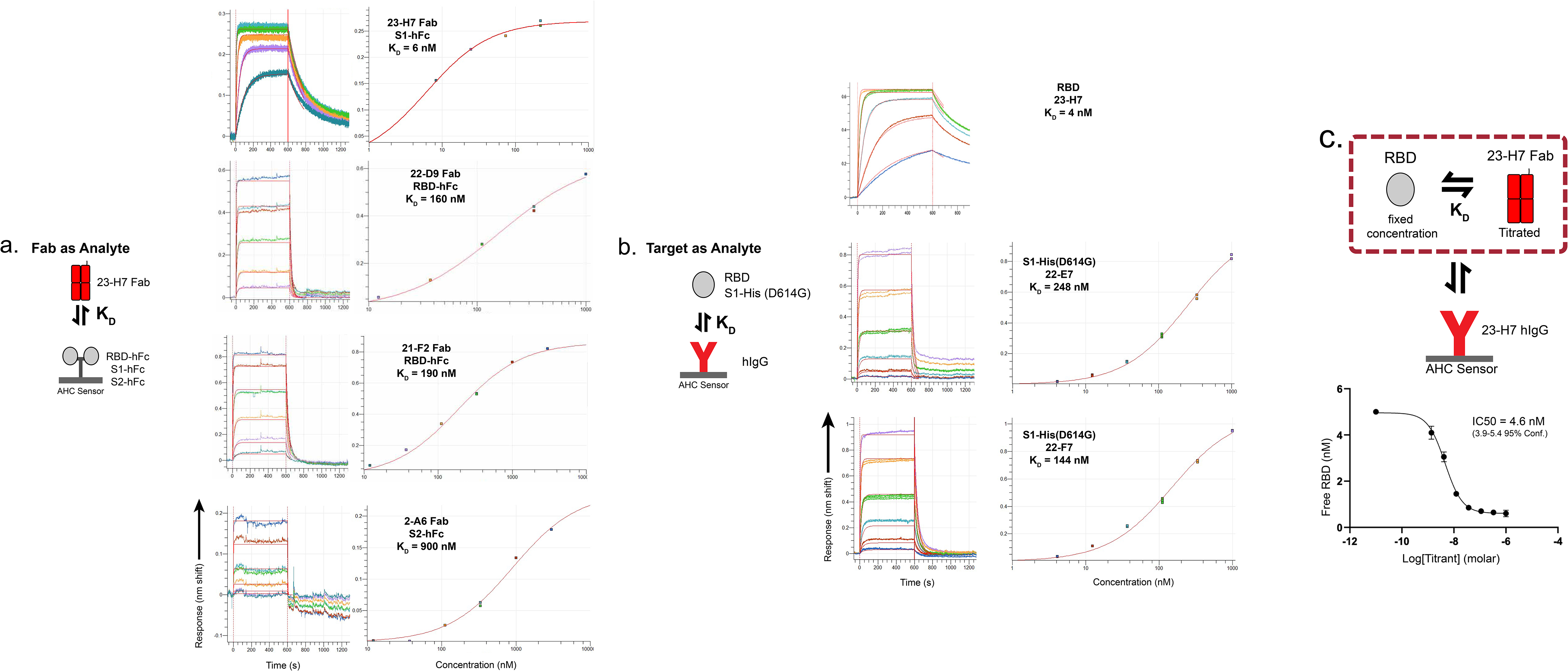
Additional graphs from multi-Ab binning assays. **(a,b)** Sensorgrams from **Fig 2a** with each binding step aligned to Y=0 demonstrating similar magnitude of signal for association of **(a)** 2-A6 (bin S2) or **(b)** 21-F2 (bin 4) regardless of whether the spike has been saturated with other Abs. **(c-e)** Sensorgrams from premix assay showing the results from sensors coated with **(c)** bin C**, (d)** ACE2-hFc, or **(e)** anti-His-mAb to probe for “free” spike binding sites in premixes of spike with various Abs, individually, or as cocktails **(see Fig 2).** Only premixes containing ACE2-blocker Abs (bin 2 or bin 4) blocked binding to ACE2-coated sensors, whereas none of the cocktails blocked anti-His-coated sensors, confirming universal access of the spike’s His tag irrespective of Ab decoration. These controls confirmed a bin-specific blockade of the spike upon saturation with Abs from up to four non-overlapping bins, simultaneously.

**Fig S4.**
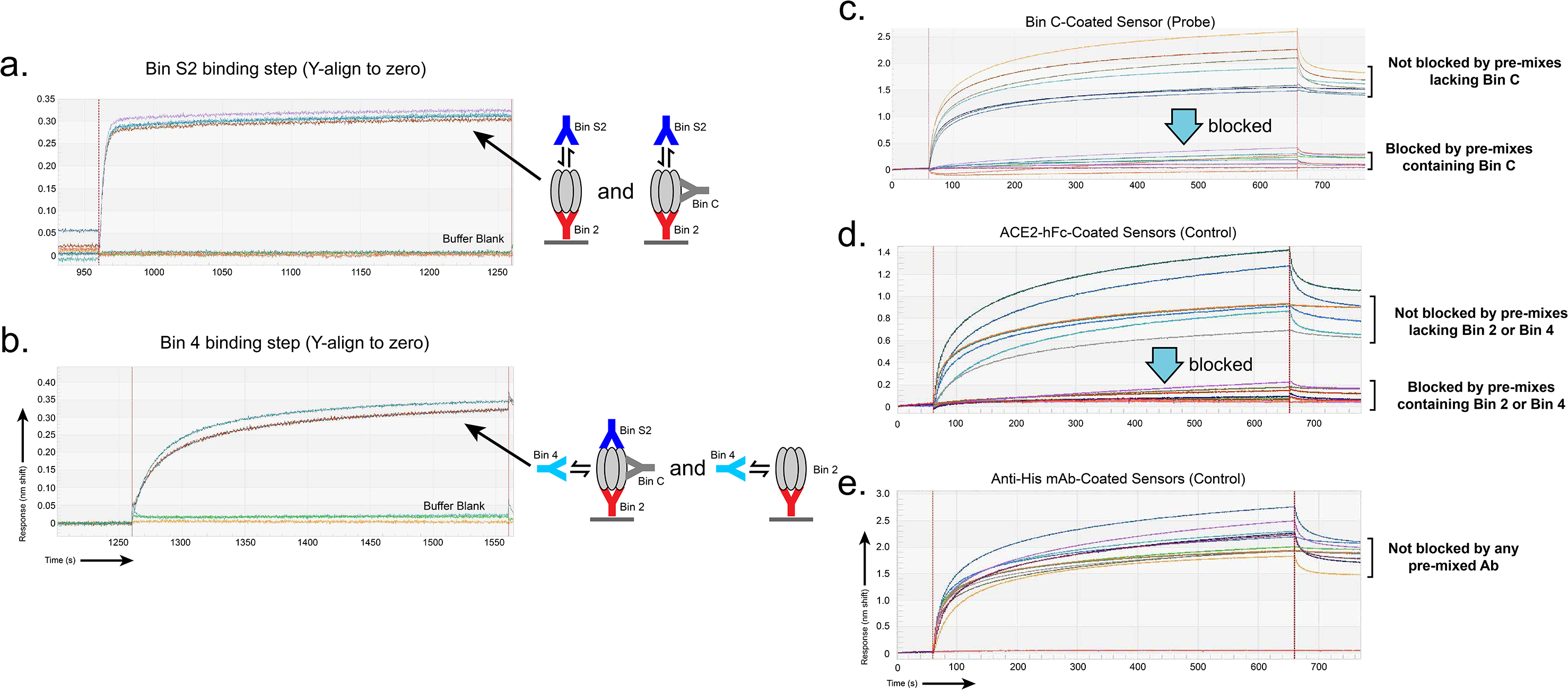
Examples of Octet-based affinity estimates. determined in complementary assay orientations, **(a)** Fab as analyte, **(b)** monovalent target as analyte and **(c)** solution affinity. The overlay plots show the sensorgrams (measured data, colored by analyte concentration) and their global fits (red lines). The kinetic (left) and steady-state (right) analysis fitting routines gave comparable affinity determinations. Summary of affinities is shown in **Supplemental Table S-1.**

**Fig S5.**
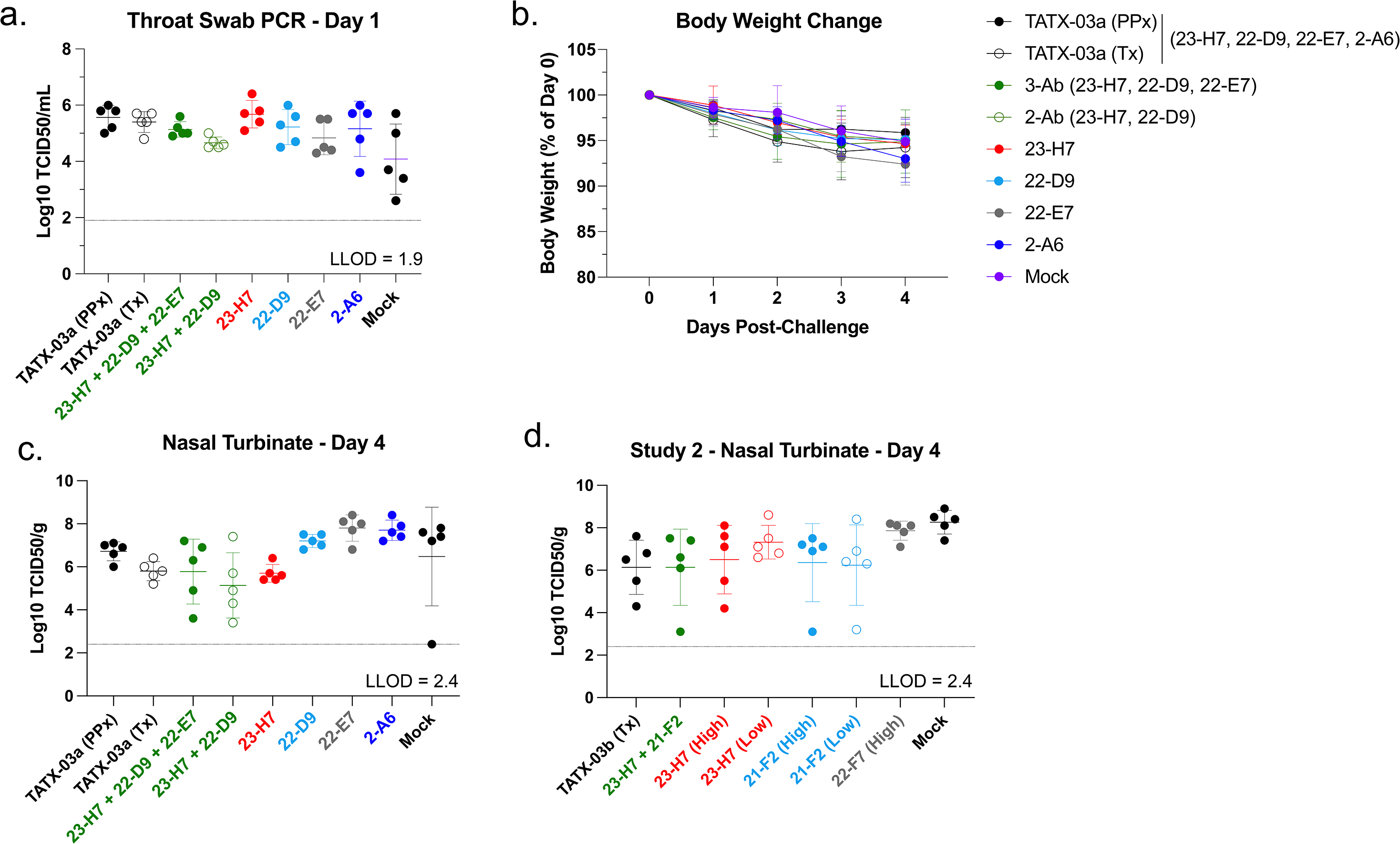
Additional data from the *in vivo* efficacy studies evaluating TATX-03. TATX-03a (23-H7, 22-D9, 22-E7, 2-A6) was administered pre-challenge (prophylactic, PPx) or post-challenge (therapeutic, Tx) as indicated at 40 mg/kg total Ab concentration (10 mg/kg/Ab in the 4-Ab blend), while the individual Abs of TATX-03a were on administered pre-challenge at 40 mg/kg. TATX-03b (23-H7, 21-F2, 22-F7, 2-A6) was administered at 20 mg/kg total Ab concentration (5 mg/kg/Ab in the 4-Ab blend) as Tx only. Individual antibodies were administered post-challenge at 20 mg/kg (23-H7, 21-F2 and 22-F7) and 5 mg/kg (23-H7 and 21-F2). In addition, one group was post-challenge treated with a 2-Ab combination of 23-H7 and 21-F2 at 5 mg/kg total Ab concentration (2.5 mg/kg/Ab). **(a)** Study 1 throat swab PCR at day 1 confirming presence of viral RNA in all animals. **(b)** Study 1 body weight change at day 4 (endpoint) expressed as a percentage of day 0 body weight **(c**-**d)** Replication-competent viral titer in nasal turbinate homogenate at endpoint for study 1 and 2, respectively. Dotted lines represent the lowest limit of detection (LLOD) for assay. See **Methods**.

**Fig S6.**
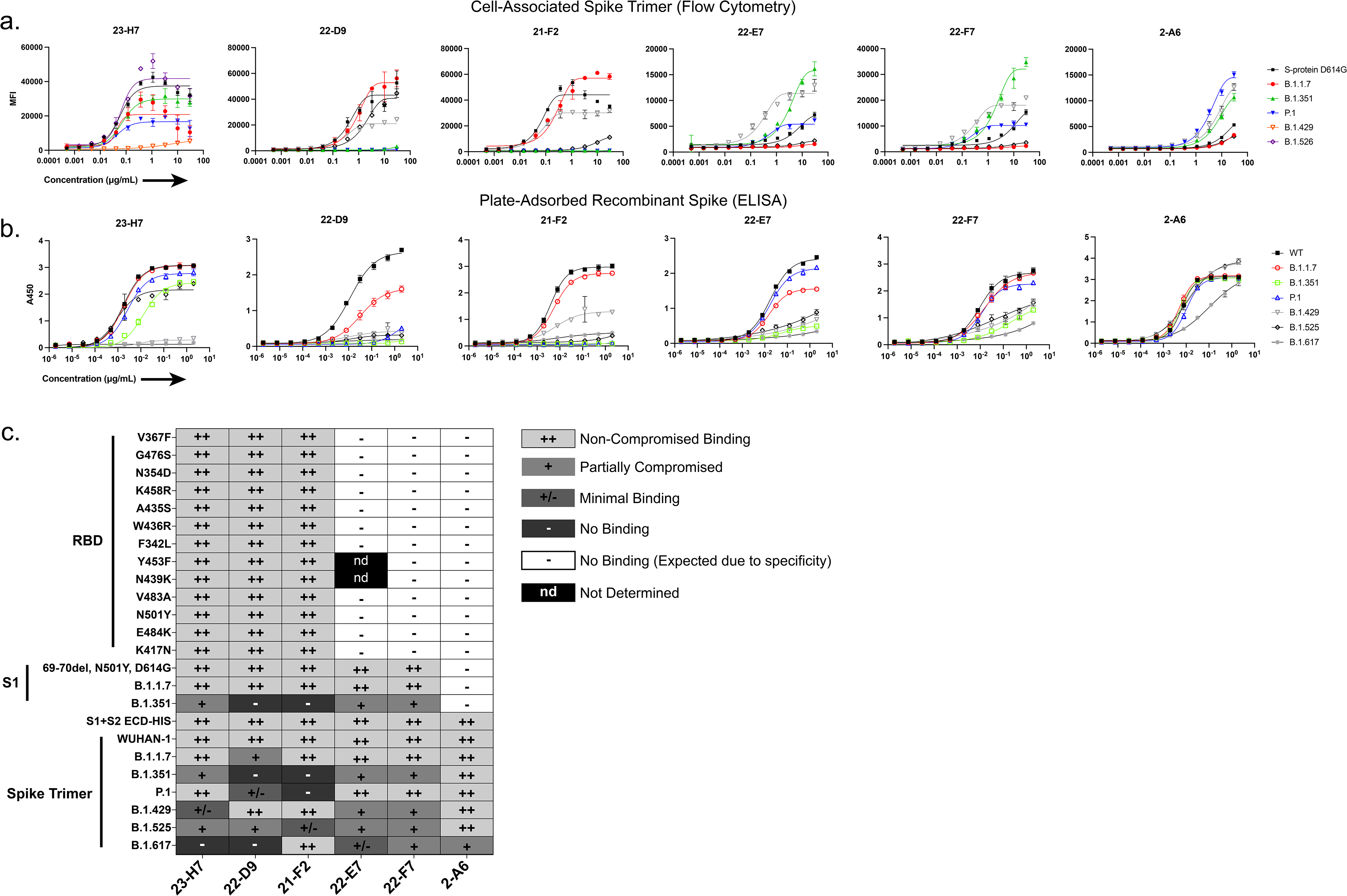
*In vitro* screening of Abs against SARS-CoV-2 variants of concern. **(a)** Binding to cell-associated spike trimer with the hallmark mutations of the specified variants. **(b)** ELISA results to plate-adsorbed recombinant spike proteins and **(c)** heat map summary of ELISA-based reactivity profiles.

**Supplemental Table-S1.**
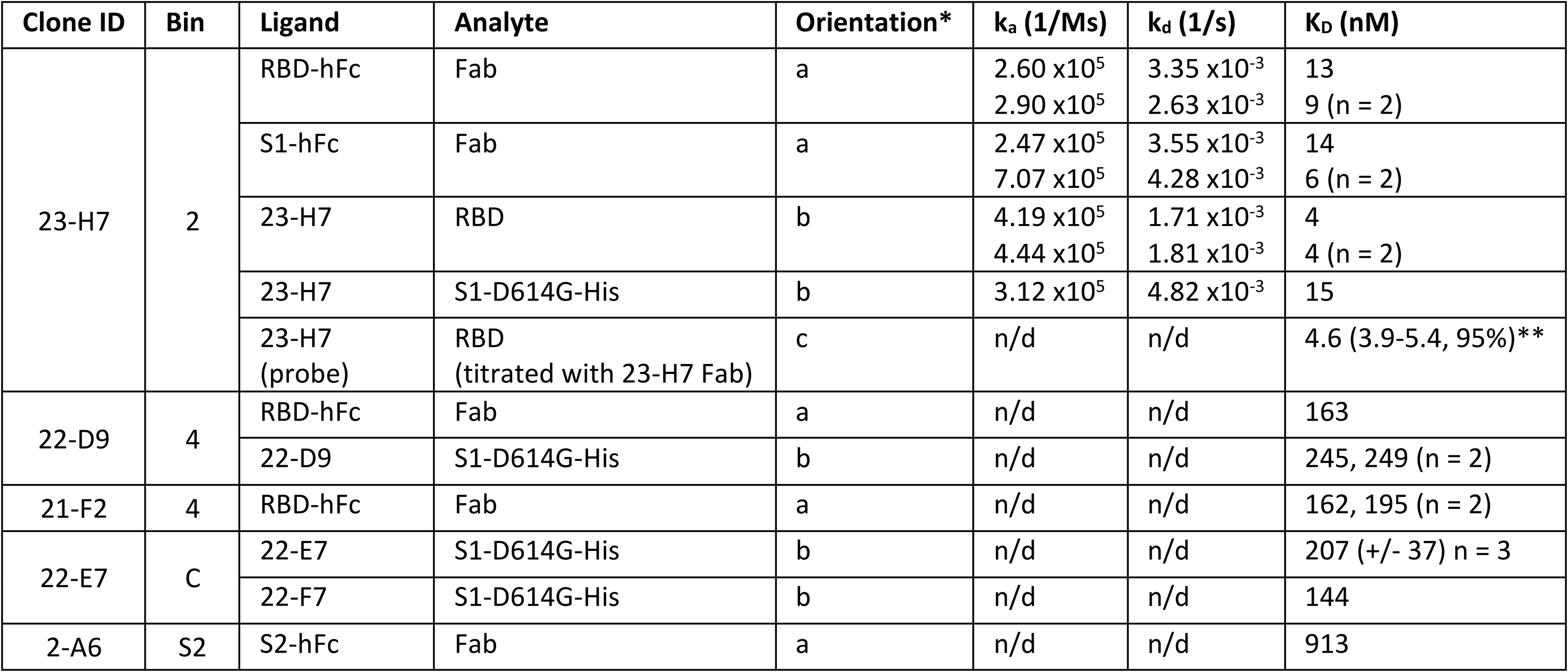
Octet affinity estimates using various assay orientations. Ligand and analyte refer to the binding partner used “on sensor” or “in solution”, respectively. n/d = kinetics not determined for steady-state analysis or solution affinity measurements *Orientations a,b,c refer to those described in Supplemental Fig S4. ** Solution affinity estimate (with 95% confidence interval)

## Notes

### Competing Interest Statement

All authors are employees of ImmunoPrecise Antibodies Ltd. which funded and carried out the studies. The authors declare no additional competing interests.

